# Hormetic curve of dietary mono- and disaccharide content determines weight gain, gut microbiota composition and cognitive ability in mice

**DOI:** 10.1101/2024.12.06.625641

**Authors:** Piotr Bartochowski, Jakub Chwastek, Bartosz Zglinicki, Olga Pietrzyk, Gabriela Olech-Kochańczyk, Monika Szewc, Aleksandra Bartelik, Julio C. Torres, Adam Karpinski, Piotr Jaholkowski, Agata Klejman, Marek Kochańczyk, Ewa Bulska, Mirosława Gałęcka, Miron Kursa, Anna Konopka, Anna Kiryk, Witold Konopka

## Abstract

Hormesis is defined as dose response phenomenon characterized by low-dose stimulation and high-dose inhibition (Calabrese & Mattson, 2017). To date, low doses of several stressors (intermittent fasting, caloric restriction or selected phytochemicals) have been shown to exert beneficial effects on health (Martin et al., 2006). In the present study, we aimed to determine hormetic factors in a series of diets used in mice. We found that animals fed high-sugar diet (HSD) or high-fat diet (HFD) containing relatively high amounts of mono- and disaccharides become obese compared to animals fed standard diet (STAND) or ketogenic diet (KD) containing low doses of these compounds. Underlying the observed metabolic phenotype may be changes in the composition of the intestinal microbiota, showing u-shaped features in selected species. It is noteworthy that a short-term dietary regimen of several weeks resulted in difficulties in achieving effective scores on a complex cognitive test based on spatial procedural acquisition in the HSD and HFD groups. Our data identify dietary mono- and disaccharide content (commonly known as sugars) as a critical hormetic factor with beneficial/harmful effects at multiple levels of body function.

## Introduction

Obesity is a pandemic of the modern world and is associated with an increased risk of life-limiting conditions such as metabolic syndrome (Eckel et al., 2005). Elevated blood pressure, insulin resistance and high triglyceride levels, all of which are strongly associated with obesity, markedly increase the likelihood of developing type II diabetes and cardiovascular disease. Two factors are commonly considered as the most likely causes of obesity: a diet with unrestricted access to palatable foods and a sedentary lifestyle, often combined with chronic stress. Their simultaneous occurrence is particularly prevalent in modern, highly developed societies (Schwartz et al., 2017). However, the straightforward nature of these mechanisms does not entirely explain the sudden rise in obesity worldwide in recent decades.

One approach to understanding the actual causes of increased susceptibility to fat deposition in adipose tissue is to compare our daily eating habits with those of our hunter-gatherer ancestors (Crittenden & Schnorr, 2017). This comparison is possible thanks to a few still existing communities that follow a diet similar to that of our ancestors from before the agricultural revolution. Despite their diverse diets based almost exclusively on wild game (Siberian populations) or plant foods (some rainforest populations), mass obesity does not occur in communities consuming solely natural goods. The diet of our ancestors contained, in varying proportions, energy sources in the form of fat and carbohydrates. The main sources of fat were hunted animals or caught fish. In contrast, plant-based foods were the main source of carbohydrates. On top of that, monosaccharides should be distinguished because of their special ability to cause rapid peaks in insulin secretion; their main sources in the primordial diet were honey and fruits (Ludwig et al., 2021). An important assumption of this study is that simultaneous consumption of fat and monosaccharides was an unlikely event in the hunter-gatherers’ diet: in all likelihood, fruit was not an accompaniment to a meal consisting of hunted game. This is in a stark contrast to the composition of modern people’s meals, where palatable food often contains large amounts of sugars and fats together. We hypothesized that the combination of these two energy sources in a meal leads to metabolic reorganization that promotes increased adiposity.

We have selected *Mus musculus* as a model organism to study the relationship between dietary composition (with particular emphasis on the ratio of fat to carbohydrate) and various aspects of body function. Mice are organisms capable of adapting to changing living conditions, and their diet can consist of both plant and animal products, the proportions of which depend on the environment and the season (Le Roux et al., 2002). This can significantly alter the composition and type of metabolically available macronutrients such as carbohydrates, fats and proteins. The mouse digestive system consists of organs that are anatomically and functionally similar to those in the human body, with some important differences, such the ratio of small and large intestines length to body mass (Hugenholtz & de Vos, 2018). Both this trait and omnivorousness make *Mus musculus* a reliable animal model for studying the effects of diet on the host body, ranging from metabolism in general, gut microbiota composition, up to cognitive function.

In this study, we observed that animals fed high-sugar diet (HSD) or high-fat diet (HFD) containing relatively high amounts of mono- and disaccharides become obese compared to animals fed standard diet (STAND) or ketogenic diet (KD) containing low doses of these compounds. Our analysis of intestinal microbiota shows that observed metabolic phenotypes are associated with changes in the composition of the microbiome. We found that short-term dietary regimen of several weeks resulted in difficulties in achieving effective scores on a complex cognitive test based on spatial procedural acquisition in the HSD and HFD groups. Overall, we identified the dietary mono- and disaccharide content as a critical hormetic factor with beneficial/harmful effects at multiple levels of body function.

## Results

### Despite equal weekly caloric intake, mice gain weight only on diets rich in sugars and fats

To investigate the effects of a diet with a varying ratio of carbohydrate to fat, the macronutrients that serve as the body’s main source of energy, we have selected four types of diets: standard chow diet (STAND), high sugar diet/Western diet (HSD), typical high-fat diet (HFD), and ketogenic diet (KD). These diets were chosen to have decreasing amount of metabolizable energy derived from carbohydrates in the order: STAND > HSD > HFD > KD, and *vice versa* for fat (Figure 1A). It is worth noting that the content of mono- and disaccharides (commonly known as sugars) was higher in the HSD and HFD diets (33.8% and 17.1%, respectively) compared to the relatively low levels in the STAND and KD diets (4.7 % and 0.7%, respectively). During the experiment, mice of C57BL/6 strain were individually housed and studied for eight consecutive weeks. The groups of mice were not significantly different in terms of body weight before experiment started. After only 1 week, mice receiving HSD and HFD diets gained significantly more weight compared to the STAND diet group (Figure 1B). This persisted for almost the entire duration of the experiment, with the exception of week 3, and by the end of the experiment (after 8 weeks), obesity was evident in the HSD and HFD groups compared to STAND group (Figure 1B,C). Mice fed all three experimental diets: HSD, HFD and KD showed reduced energy expenditure during the dark/active phase compared to STAND-fed mice (Figure 1D). This shift may partially explain the metabolic changes that induced elevated fat accumulation in mice on HSD and HFD diets. Consistent with circadian rhythms, mice had reduced energy expenditure during the light/rest phase. However, this effect was not observed in KD mice, which may account for the lack of overall weight gain in this group (Figure 1D).

**Figure 1.**
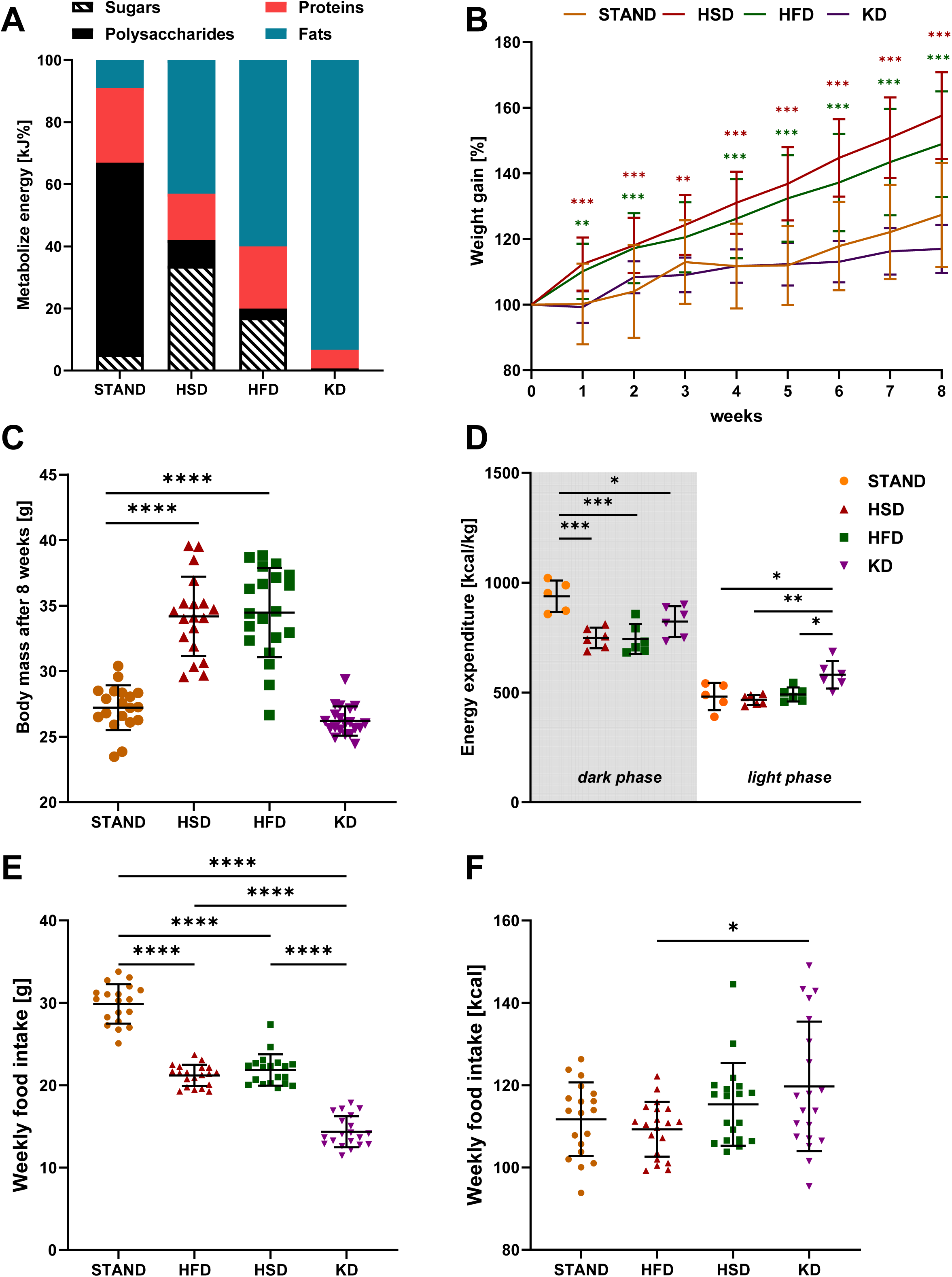
Body weight, energy expenditure and weekly food intake according to diet type. **A,** Composition of diets in terms of metabolisable energy used in the experiment according to the manufacturer’s description presented as % of energy provided by each macronutrient (fats, polysaccharides, proteins and sugars). **B,** Weekly diet-dependent relative weight gain of animals, calculated as a percentage of the weight of each animal in week 0 (beginning of the experiment) (n=19-20, mean ± SD). **C,** Absolute body weight of animals after 8 weeks of feeding individual diets (n=19-20, mean ± SD). **D,** Energy expenditure of animals related to the diet, in the dark and light phases, calculated as kcal per kg of body weight (n=5-6, mean ± SD), **E and F,** Average weekly food intake presented as weight or calories of food consumed, respectively (n=19-20, mean ± SD). \**p* ≤ 0.05, \*\**p* ≤ 0.01, \*\*\**p* ≤ 0.001, \*\*\*\**p* ≤ 0.0001; one-way ANOVA Tukey’s post hoc test, except B (One-way ANOVA with Dunnett’s multiple comparison test comparing each time point to week 0, the color of the asterisks is appropriate to the type of diet).

Further explanation of the obesity phenotype observed in HSD and HFD mice can be provided by analysis of food intake. After two weeks of adaptation to the new diet, during which the mice’s daily food intake stabilised, there were significant differences in food consumption between diets (STAND > HSD = HFD > KD, in grams) (Figure 1E). Most surprisingly, given the different caloric density of the fodders, total weekly caloric intake was similar in each dietary group (Figure 1F), and yet it lead to different weight gains.

### Diet-dependent alteration of the composition of the mouse gut microbiota

Previous studies indicate that dietary profiles can alter the gut microbiota (Tuck et al., 2020). Our data showed that total bacterial abundance had increasing trend in all three experimental diets (HSD, HFD, KD) compared to the control group (STAND) (Figure 2A), moreover, we did not observe any significant changes in species diversity, only a reduction trend in mice fed the HSD diet (Figure 2B). Furthermore, significant changes in the ratio of *Bacteroidetes* to *Firmicutes* were found. While the ratio was similar in mice fed the STAND and KD diets, it differed significantly in groups fed the HSD and HFD, but in the opposite directions (Figure 2C). These phenomena were observed to be due to the total abundance of the *Bacteroidetes* phylum in a diet rich in sugar and fat (Supplementary Figure 1B). Of note, in the case of the HFD group, almost complete dominance of the phylum *Firmicutes* in the composition of fecal microbiota was observed (Figure 2C and Supplementary Figure 1B). A more detailed analysis showed that the diversity of bacterial species belonging to the phylum *Bacteroidetes* has an increasing trend in HSD and HFD compared to STAND and KD (Supplementary Figure 1A), which was not observed for the *Firmicutes* phylum (Supplementary Figure 1A). Although global change in the abundance of major bacterial divisions cannot explain fully observed changes in metabolism homeostasis resulting in weight gain of HSD and HFD mice, we have identified individual species potentially contributing to this phenotype by applying the Boruta feature selection algorithm (Kursa & Rudnicki, 2010) (Supplementary Figure 2). The method is capable of identifying multivariate and non-linear interactions, including the hormetic (U-shaped) ones. In order to detect bacterial species whose abundance exhibits hormetic characteristics in obese compared to non-obese animals (U-shaped or inverted U-shaped), the data were further sorted according to the weight gain of individual mice (from lowest to highest), resulting in clustering of animals according to diet consumed (Figure 2D). The results showed hormetic changes in the abundance of bacterial species in the phylum *Firmicutes* (19 species) and seven species belonging to the phylum *Bacteroides* (from families *Bacteroidaceae, Prevotellaceae* and *Rikenellaceae*). The family *Lachnospiraceae* (*Firmicutes* cluster) appeared to be the most affected by diet, with 13 species significantly changed abundance, including 9 species down-regulated in mice fed HFD and HSD diets (Figure 2D).

**Figure 2.**
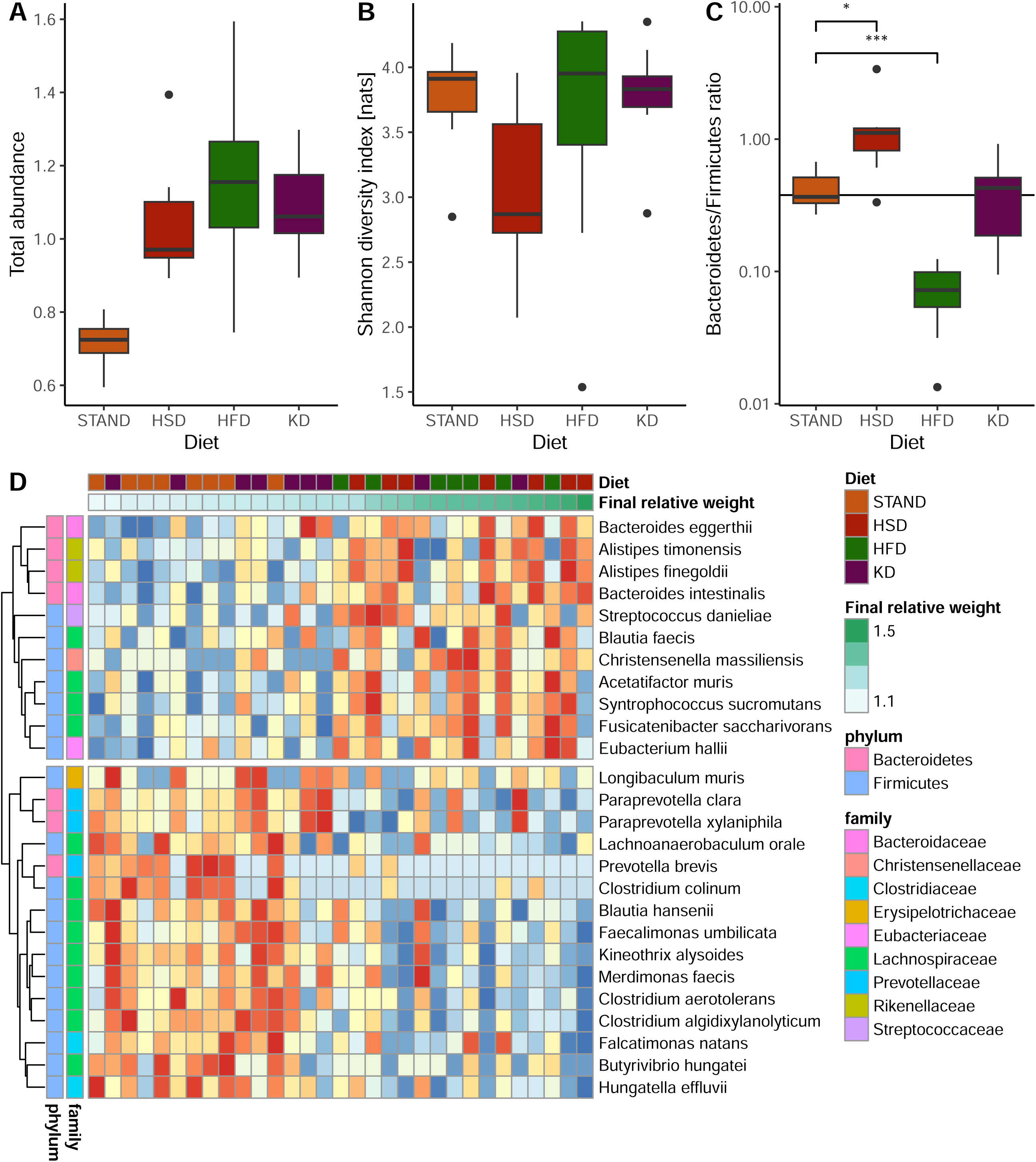
16S rRNA analysis of the diet-associated faecal microbiome. **A,** Total abundance of bacteria, **B,** Species diversity according to Shannon’s method and **C,** Bacteriodes/Firmicutes phyla abundance ratio, all in relation to diet. **D,** Heatmap showing abundance of species presenting hormetic traits, in relation to final weight gain (at the end of the experiment), according to the machine learning analysis (Kursa & Rudnicki, 2010). Abundances are shown as a ranking of all mice, with blue indicating the lowest and red the highest abundance.

### Diet composition significantly modulates mouse metabolism-related blood parameters

The results of the blood serum analysis showed a strong effect of the diet on the metabolism of the mice. The fasting blood glucose levels of the mice did not differ between the dietary groups. In non-fasted animals, a significant change in this parameter was only observed between the HFD and KD groups (Figure 3A). Mice in the HFD and KD groups showed elevated levels of free fatty acids, while the main ketone body, β-hydroxybutyrate (BOH), was elevated in the blood of KD animals, indicating a corresponding state of ketosis (Figure 3B,C). GC-MS analysis of small molecules in blood samples made it possible to distinguish samples according to the type of diet used, as shown by principal component analysis (PCA) plot (Supplementary Figure 3). Elevated levels of certain compounds were due to the feed composition, such as substances of plant origin (STAND), saccharose (HFD or HSD), cholesterol (HFD and KD) or saturated fatty acids (HFD, HSD and KD) (Figure 3D). Interestingly, animals fed HSD and HFD showed increased levels of oleic acid (monounsaturated acid, MUFA), while the palmitoleic acid (MUFA) was higher only in the HSD group. On the other hand, linoleic acid (polyunsaturated fatty acids, PUFA) level was elevated in the plasma of mice on the STAND and KD diets. Differences in saccharide metabolism were also observed, with high plasma levels of glucopyranose and ribose in the control group, while arabinofuranose and fructose were present in the other diets. In addition, aceturic acid was present at high levels in animals that were not obese and was absent in groups with weight gain. Notably, serotonin, which was present in the other groups, was not detected in the plasma of KD-fed mice (Figure 3D).

**Figure 3.**
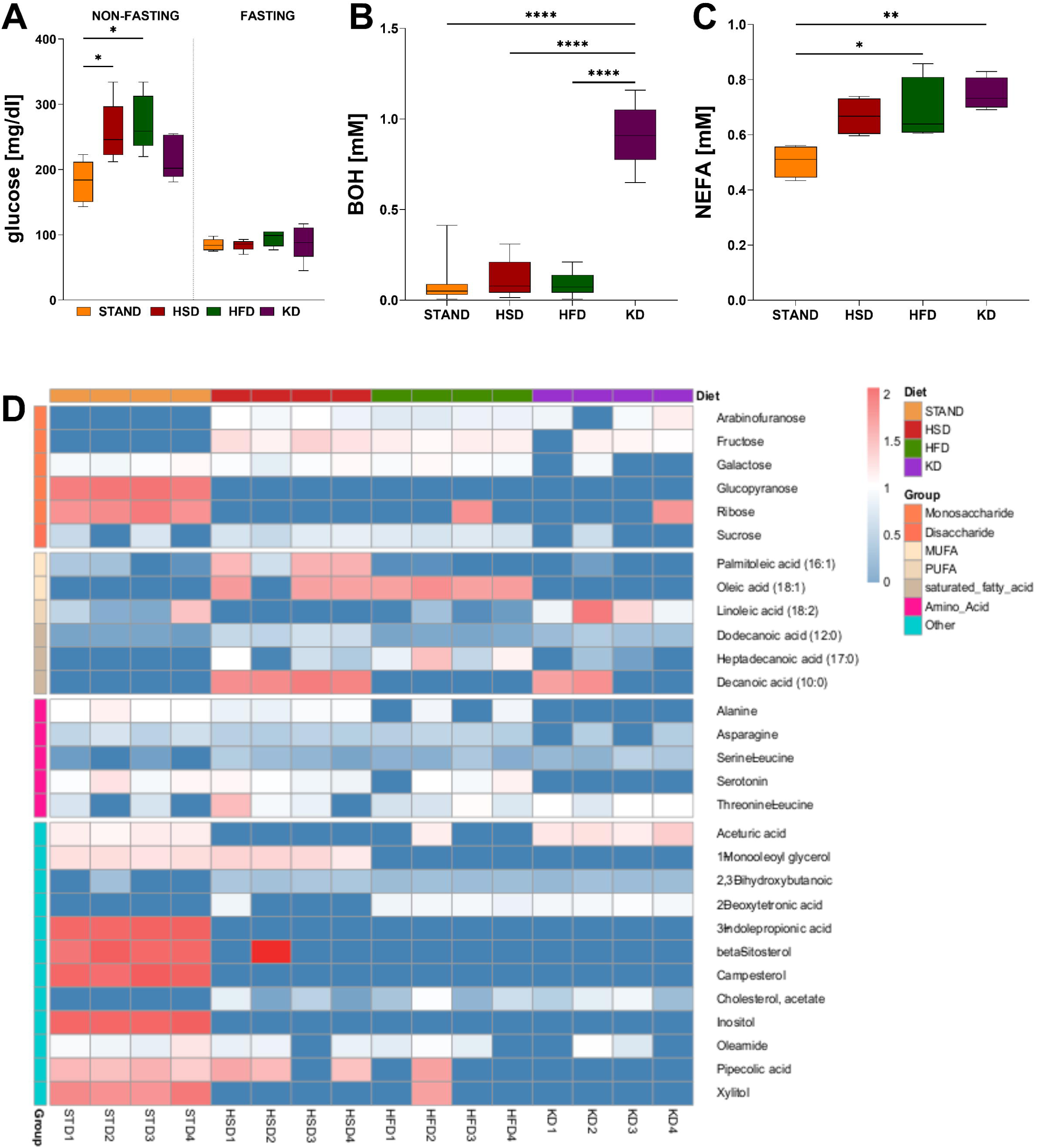
Diet-dependent modulation of metabolic parameters in mouse plasma. **A-C,** Plasma levels of primary metabolites such as glucose (n=5), beta-hydroxybutyrate (BOH) (n=10-13) and non-esterified fatty acids (NEFA) (n=4). Data are presented as box plots ± min/max value and analyzed using one-way ANOVA with Tukey’s post hoc test (\**p* ≤ 0.05, \*\**p* ≤ 0.01, \*\*\**p* ≤ 0.001, \*\*\*\**p*≤ 0.0001). **D,** Normalized GC-MS data (n=4) are presented as a heatmap, sorted by type of diet and group of compounds analyzed.

### Diets modify blood circulating miRNAs profile

MicroRNA profiling in plasma showed that diets impacted the composition of these molecules, which are postulated to mediate brain-gut communication (Ferrero et al., 2020). From the study pool, we selected nine microRNAs that differentiated the dietary groups according to the phenotypic changes observed (Figure 4). Significant differences in plasma miRNA abundance were observed between STAND and both obesity-inducing diets (mmu-miR-101a; mmu-miR-101b; mmu-miR-122; mmu-miR-192; mmu-miR-194; rno-miR-345-3p), while two of them were also overexpressed in KD relative to STAND (mmu-miR-101b; mmu-miR-194). On the other hand, comparison of the expression of the analyzed miRNAs versus KD showed a significant difference between HFD (mmu-miR-101a; mmu-miR-101b; mmu-miR-122; mmu-miR-148a; mmu-miR-192; mmu-miR-500; mmu-miR-802), and only rno-miR-345-3p increased both in HFD and HSD vs. KD. Meanwhile, the expression levels of three other miRNAs (mmu-miR-101a; mmu-miR-101b; mmu-miR-148a) were significantly higher in the HFD group compared to HSD (Figure 4).

**Figure 4.**
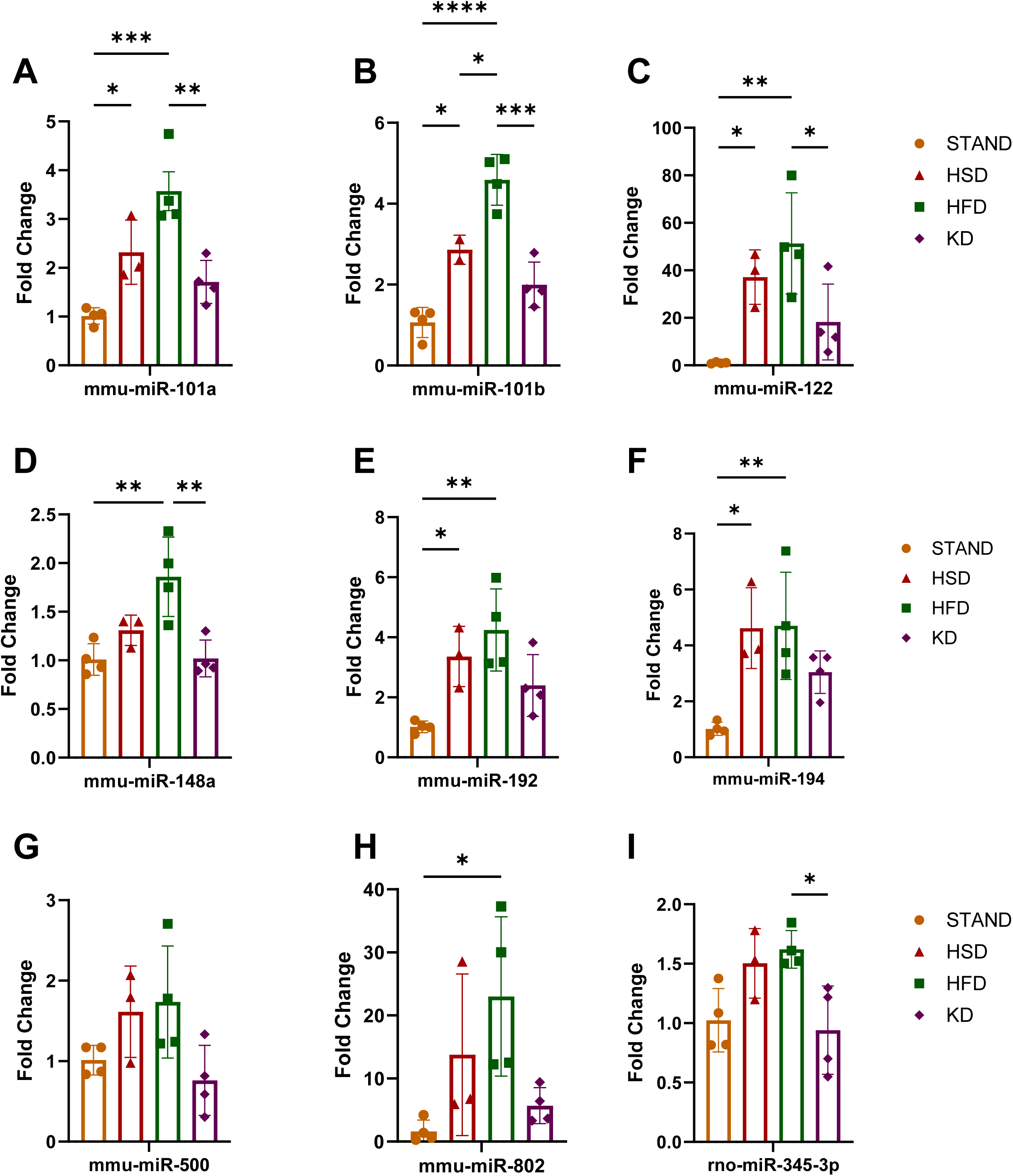
Hormetic characteristics of selected miRNAs in mouse plasma. Relative changes of miRNA (**A,** mmu-miR-101a, **B,** mmu-miR-101b, **C,** mmu-miR-122, **D,** mmu-miR-148a, **E,** mmu-miR-192, **F,** mmu-miR-194, **G,** mmu-miR-500, **H,** mmu-miR-802, and **I,** rno-miR-345-3p) expression in plasma of mice fed different diets. Data were normalized to global miRNA expression and calculated as fold change in expression of each analyzed miRNA in the STAND group (STAND, HFD, KD – n=4; HSD – n=3). \**p* ≤ 0.05, \*\**p* ≤ 0.01, \*\*\**p* ≤ 0.001, \*\*\*\**p* ≤ 0.0001; one-way ANOVA with Tukey’s post hoc test.

### The cognitive functions of mice fed HSD and HFD diets are impaired

Dietary changes cause direct or indirect (*via* gut microbiota-derived metabolites) effect on the host metabolism. It is well known, that the overall metabolic state of the body influences the cognitive ability through numerous mechanisms (van Praag et al., 2014). Cognitive function of mice fed all four diets for 8 weeks (STAND, HSD, HFD, KD) was assessed under home cage conditions in IntelliCage (Kiryk et al., 2020). In order not to interfere with the diet, and thus the metabolism of the mice, or the composition of the microbiota, we designed a non-aversive, non-appetitive (without using sweetened water) complex procedural test based on spatial memory. To drink plain water, mice were forced to visit corners in a specific order, either clockwise or counterclockwise, each corner in turn (Thirst-based cognitive test, Figure 5A,B). For the reference animals (unpublished preliminary experiments and STAND animals), it took about 9 days to learn and stabilize the rule of alternating the corner after each drinking session (to the next one on the right) and develop a stable response rate of more than 60% correct visits (Figure 5C). The learning curve was plotted by a decreasing number of incorrect visits, while the number of correct visits remained relatively constant, thus expressing the level of thirst (Supplementary Figure 4). Not only was rule memory required, but also spatial memory to remember the location of the next corner when selecting it. Choosing the wrong corner resulted in a non-opening door and a lack of access to water. A single return to the current corner was considered neutral (not treated as an incorrect visit) and mice made a minimal number of such return visits (Supplementary Figure 4). Thus, based on the probability level, the success rate in choosing the right corner should be about 33%. All animal groups tested achieved this level of correct visits on the first day of the experiment (Figure 5C). The performance of the mice in the thirst-based cognitive test was influenced by the type of diet they received. STAND and KD mice started to acquire the drinking rule as early as day 2 of the experiment, and by day 3, more than 50% of visits were made at the correct corner (Figure 5E). In contrast, HSD and HFD mice began to improve only on day 4 or 5 and did not reach 50% correct responses until day 7. Eventually, all groups reached a similar level of correct responses. The first part of the experiment was followed by a re-learning test, in which the mice had to perform a sequence of visits and nosepokes in a counter-clockwise manner (Figure 5B). In this case, the reduced performance of the HSD and HFD groups was observed only on day 1 after the change in experimental regimen (Figure 5F). Although the HFD group slightly improved its performance (Figure 5D), there were no significant differences between the groups on the other days (Figure 5F). To sum up, we show that HSD and HFD diets can negatively affect the learning of complex cognitive tasks.

**Figure 5.**
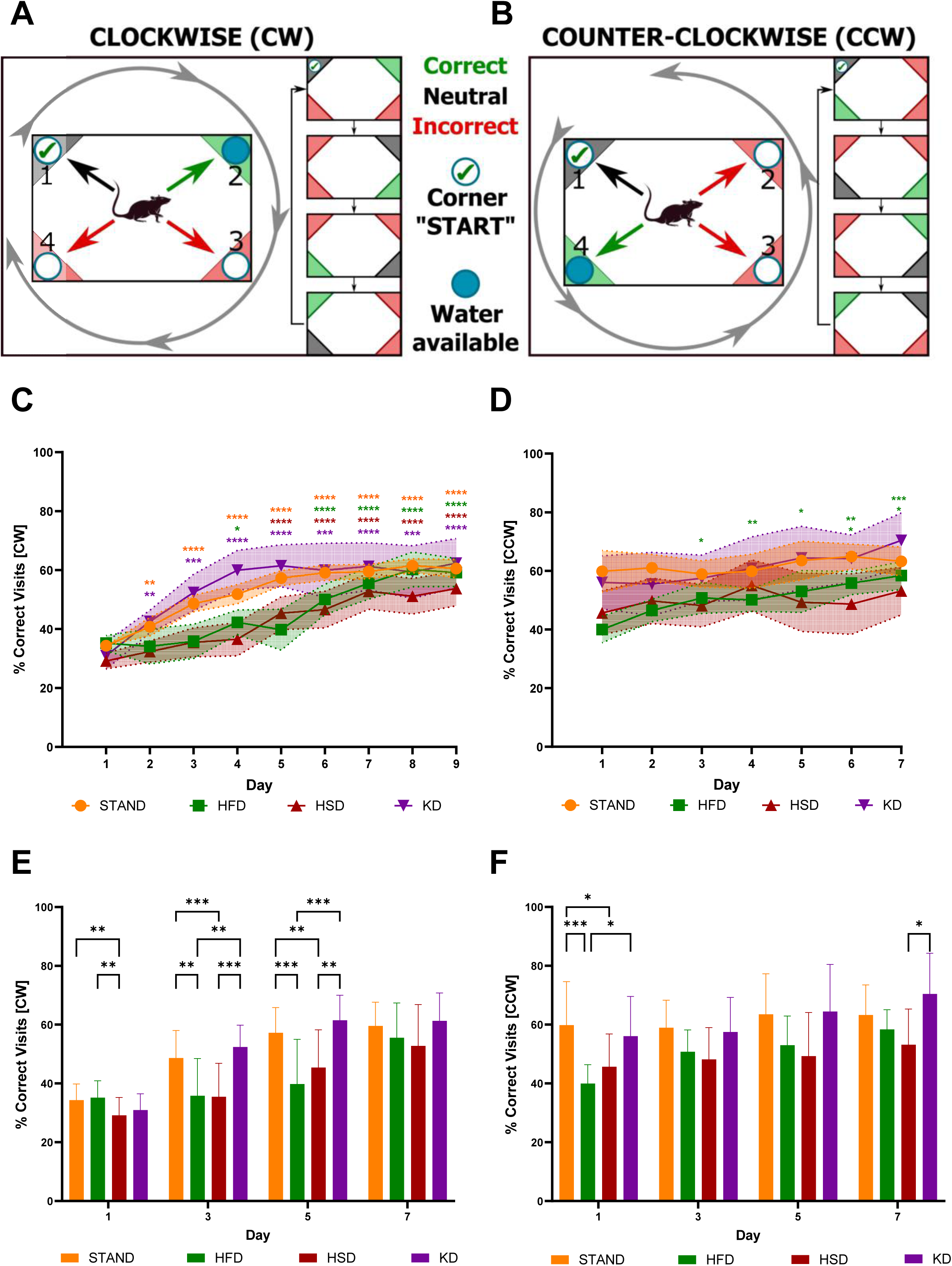
Impaired cognitive functions of mice fed HSD and HFD diets. **A and B**, Graphic diagram of clockwise (CW) and counterclockwise (CCW) behaviour test. In the CW test, the animal, after first selecting a corner with access to water, had to go clockwise (to the right corner) to drink again (correctly), returning to the corner was not counted as an error (neutral, no access to water), but selecting one of the other two corners was incorrect (no access to water). During the CCW test, the principle was the same, but the animal moved counterclockwise (to the left corner). **C and D,** The learning curve expressed as a percentage of correct visits in the CW and CCW tests, respectively, the data from each time point was compared to the first day of the experiment for each diet. Experiments were conducted for 9 (CW) or 7 (CCW) days((CW n: STAND = 46; HFD = 21; HSD = 23; KD = 8); (CCW n: STAND = 19; HFD = 10; HSD = 11; KD = 8), mean ± 95% CI). **E and F,** Percentage of correct CW and CCW visits on days 1, 3, 5, and 7 from the start of testing and a comparison of the differences in this parameter by diet at each time point (mean ± SD). \**P* ≤ 0.05, \*\**P* ≤ 0.01, \*\*\**P* ≤ 0.001, \*\*\*\**P* ≤ 0.0001; one-way ANOVA with Dunnett’s (**C and D**) (comparing each time point to day 1, the color of the asterisks is appropriate to the type of diet) or two-way ANOVA with Tukey’s (**E and F**) post hoc test.

## Discussion

Hormesis is generally considered an adaptive response of the biological system to low-dose challenging stimulation, as originally defined for ionizing radiation (Luckey, 2006). Of note, our analysis of dietary composition showed that mono- and disaccharide content alone (low in STAND and KD, high in HSD and HFD) has hormetic effect on the development of obesity, composition of gut microbiota and finally cognitive function.

To date, obesity has most often been associated with excessive dietary calorie intake (Wright & Aronne, 2012). In addition to the many fashionable dietary regimes in developed countries, diets rich in sugar and fat, known as Western diets, are common and lead to the development of obesity (Clemente-Suarez et al., 2023). Our experiments confirmed that animals fed such diets gained significantly more weight compared to the control (chow diet) and ketogenic diet groups. Despite the different diet composition and the observed increase in adiposity in the HSD and HFD groups, the amount of calories consumed by studied mice were not different across groups. Therefore, the obesity in the HSD and HFD groups may not be due to increased consumption and deposition of excess energy. An alternative view of the metabolic cascade triggered after eating a meal (especially one containing high amounts of mono- and disaccharides) is described by the carbohydrate-insulin model (CIM) (Ludwig & Ebbeling, 2018). According to this theory, food with a high glycemic index (mostly containing sugars) triggers an unusually high insulin secretion. This primarily stimulates anabolic processes such as fat storage in adipose tissue. At the same time, levels of circulating energy sources (glucose, lipids) are reduced, which is sensed by the brain, able to precisely control the satiety-hunger balance (Sternson & Eiselt, 2017; Vinnikov et al., 2014), resulting in reduced energy expenditure and, paradoxically, increased feeling of hunger. When a meal combines sugars and lipids, fat storage is facilitated. In animal models, insulin administration promotes fat deposition and a reduction in energy expenditure (Cusin et al., 1992) and diets containing high glycemic load foods induce fat deposition (Pawlak et al., 2004). The effect of hyperinsulinaemia is reflected in patients with type II diabetes, in whom weight gain is a common side effect of insulin therapy (Carlson & Campbell, 1993). The carbohydrate-insulin model should not be considered as the opposite to the conventional caloric balance model, but as its extension emphasising the importance of a qualitative approach to calories uptake. The carbohydrate-insulin model correctly reflects the correlation of the glycemic load of the studied diets with the development of obesity. The HFD and HSD diets had a high glycemic load due to their high sugars content. This, coupled with high amounts of saturated fats, cause excessive body fat growth. According to CIM, the lack/low level of carbohydrate in the KD diet does not induce an insulin peak, preventing fat sequestration in adipose tissue and inducing increased energy expenditure (Hall et al., 2016). Such an effect observed in the resting phase protected KD mice from developing obesity. On the other hand, the STAND diet contains mostly complex carbohydrates with a lower glycemic index and is low in fat.

Consistent with the carbohydrate-insulin model (in which the total flow rate of a fuel such as glucose is important, rather than its instantaneous concentration), we detected no differences in baseline glucose measurements, except for a slight increase in the HFD group. Similarly, we observed a normal fasting glucose response in all groups, indicating the absence of type II diabetes in these mice. It is likely that the too short duration of the dietary intervention (8 weeks) only resulted in the development of obesity without co-morbidities (Stott & Marino, 2020; Surwit et al., 1988). Our extended metabolomic analysis revealed two compounds exhibiting a hormetic factor pattern: N-acetylglycine (aceturic acid) and oleic acid. For the latter, in line with our results on HFD and HSD diets, it was shown that obese animals on high-fat and high-sugar diets showed increased triglycerides concentrations containing oleic acid derived from lipogenesis in the liver (Sato et al., 2010). Changes in the metabolites circulating in the blood may originate from diet itself or be products of host and gut microbiota metabolism. As an example, we observed significantly higher plasma levels of aceturic acid in control and KD-fed animals compared to the obese groups, which is particularly interesting given previous studies in which supplementation of HFD-fed mice with this compound inhibited weight gain (Fluhr et al., 2021). Moreover, human studies have shown that aceturic acid serum concentration was inversely associated with the risk of metabolic syndrome development (Perng et al., 2017). However, the precise composition of bacterial species modulating the concentration of N-acetylglycine is unknown (Shapiro et al., 2022).

Diet strongly affects the composition of the gut microbiota, which is reflected in the host at the metabolomic and proteomic level (Daniel et al., 2014; David et al., 2014). Recent accumulating evidence suggests that the gut microbiota plays an important role in the development of obesity (Turnbaugh, Baeckhed, et al., 2008). In particular, on the one hand, weight gain can be induced by colonizing germ-free mice with the obesity-associated gut microbiota phenotype (Turnbaugh et al., 2006), while on the other hand, beneficial bacteria can be used to reverse diet-induced obesity (Kong et al., 2019). In a healthy organism, the gut microbiota remains in a state of eubiosis, in which a highly biodiverse and balanced microbial ecosystem benefits the host (Hollister et al., 2014). One of the hallmarks of the composition of the gut microbiota associated with obesity is a disruption in harmonious composition, leading to a disruption in mutualistic interaction with the host known as dysbiosis (Turnbaugh, Backhed, et al., 2008). Research findings, including the data we present here, point in different directions of these changes (Clarke et al., 2012), making it difficult to use this indicator as an unequivocal biomarker of obesity and other pathological conditions. There may be multiple explanations for these inconsistencies, including the influence of host genetics, environmental factors (including diet), as well as timing of sampling and lack of standardisation of microbiota testing (Jarett et al., 2021; Sergaki et al., 2022; Sinha et al., 2017).

In our study, the bacterial family, whose abundance varied in a hormetic trend according to the sugar content of the diet used, was most frequently represented by the *Lachnospiraceae*, making this group potentially key to the substrate development of the changes observed in the experiment. Under these assumptions, high levels of bacteria from this family may be beneficial due to their ability to produce SCFAs (Short-Chain Fatty Acids) such as acetate, butyrate and propionate (Biddle, 2013; Sheridan et al., 2016). Thus, for example, we observed increased abundance of *Blautia hansenii* in animals fed STAND and KD. The species has previously been shown to negatively correlate with visceral fat and insulin levels in a sex-dependent manner (Hases et al., 2023). Accordingly, *B. hansenii* supplementation reduced white adipose tissue growth and improved glucose tolerance in mice treated with a high-fat diet (Shibata et al., 2023). Likewise, in humans it was shown that two species: *B. hansenii* and *B. producta* were negatively correlated with visceral adipose tissue area (Ozato et al., 2022).

The effect on cognition may be through diet-induced altered metabolic homeostasis sensed by the hypothalamus (Ramirez et al., 2022). Numerous studies also indicate that obesity is associated with cognitive impairment, often leading to neurodegenerative processes (Andre et al., 2019; Nguyen et al., 2014). Although the mechanisms underlying the impact of a high-fat diet on neurodegenerative processes have not yet been established, the literature argues that neuroinflammation may be the factor accelerating this process (Castanon et al., 2015). Additionally, gut microorganisms can remotely/indirectly influence the brain through substances they produce, such as SCFA or bile acids (Mulak, 2021; Torres-Fuentes et al., 2017). Anxiety-like behaviour or short-term working memory may be impaired by gut-derived metabolites in both humans and mice (Arnoriaga-Rodriguez et al., 2020; Needham et al., 2022).

To reliably detect subtle cognitive changes in mice fed different diets (with no apparent pathological symptoms), we used a complex test that took the animals several days to solve it. This procedure represents a different approach from that used by other researchers, where mice solve tasks in sessions lasting a few minutes or at most a few hours. Assessment of cognitive function with IntelliCage system additionally allowed us to avoid stress-inducing conditions that could affect the microbiome, such as aversive arenas, social isolation or human contact. Similar working memory tests have been used in IntelliCage previously by others (reviewed by (Kiryk et al., 2020)). However, the researchers typically applied drinking sessions (for example, 2 sessions of 1 hour of drinking per 24 hours), which can present additional difficulty for mice if they do not know when a drinking session starts and ends. In our study, the mice were allowed to drink and learn all the time. At the same time, we limited the drinking time per visit to 7 seconds so that mice could not drink at will in one corner. We believe that this approach allowed us to better study the dynamics of spontaneous learning based on water reward motivation. In addition, we used a light stimulus that indicated the possibility of a reward at a particular corner. We also excluded tests involving aversive stimuli (air-puff or electric shock) or appetitive reinforcement (sweet water/food) that would directly alter diet composition. Instead, we have applied a complex, procedural cognitive test during home cage monitoring as discussed before (Kiryk et al., 2020). The test exploits the natural tendency of mice to patrol all corners of the cage which was shown to be hippocampus-dependent (Voikar et al., 2010).

MicroRNA-dependent regulation of physiological processes, in particular neurophysiological processes, can be considered at several levels. The primary action of microRNAs is to repress translation of target mRNAs, a phenomenon observed in both plants and animals (Bartel, 2009). Inside the neuron, these molecules directly affect neuronal function by regulating the translation of target genes associated with neuronal excitability and plasticity (Fiorenza & Barco, 2016). As we and others have shown, global deletion of microRNAs from mouse brain neurons has demonstrated their role in regulating learning and memory (hippocampus and cortex dependent) (Fiorenza et al., 2016; Konopka et al., 2010), as well as food intake behaviour (hypothalamus dependent) (Fiorenza et al., 2016; Mang et al., 2015; Vinnikov et al., 2014). Active microRNAs can be of endogenous, intracellular origin or derived from other cells or tissues. These so-called circulating microRNAs are transported by the blood as extracellular vesicles or complexed with proteins, allowing them to regulate functions in distant tissues (Mori et al., 2019). They are secreted by various tissues, including metabolically relevant ones such as adipose tissue (Ji & Guo, 2019). We identified nine miRNAs whose levels were elevated in animals fed an obesogenic diets, but not ketogenic or standard diet. In relation to the metabolic syndrome, serum levels of miR-101 and miR-802 were elevated in patients with type 2 diabetes (Higuchi et al., 2015). In addition, miR-122, miR-192 and miR-194 were elevated in the plasma of overweight/obese patients in the insulin resistance group (Shah et al., 2017). Interestingly, miR-148 has been found to be involved in the regulation of the inflammatory response (Friedrich et al., 2017), which is chronically increased in obese individuals. In the context of the differences in cognitive function found in mice fed different diets, it seems interesting that the researchers showed that miR-148 expression levels, among others, were significantly increased in the blood of mice with age-dependent cognitive impairment (Islam et al., 2021). When the primary hippocampal neurons culture was supplemented with miR-148 and miR-181a, a decrease in the expression of genes responsible for synaptic function and neuronal plasticity was observed. In addition, a reduction in the number of dendritic spines and abnormal neuronal firing activity in these neurons was also observed (Islam et al., 2021). In light of the limited data above, the relationship between miRNAs, metabolism and cognitive function seems to be an interesting direction that requires further research.

The main finding of this work, demonstrating that the mono- and disaccharide content of the diet acts as a hormetic agent, may imply that manipulating only this select group of compounds in the diet may have a desirable therapeutic effect against obesity, neurodegenerative diseases or ageing. In this context, low-sugar standard and ketogenic diets can already be considered low-dose stimulation compared to high-sugar Western diets, without the need to resort to more radical interventions like caloric restriction or intermittent fasting (Martin et al., 2006).

## Methods

### Animals and study design

Male C57BL/6 mice were obtained from Janvier Labs and housed individually in IVC cages GM500 green line supported by Smart Flow AHU designed by Tecniplast under conditions of 22±2 °C temperature and 55±15% humidity with *ad libitum* feeding of a standard diet, under a 12-h light/dark cycle. After 2 weeks, mice were divided into 4 groups: group fed a standard diet (STAND, 3.23 kcal/g; Ssniff v1534-703) consisting of 67% carbohydrates, 24% proteins, and 9% fat; group fed a high sugar high-fat diet (HSD, 4.60 kcal/g, Ssniff E15775-34) consisting of 42% carbohydrates, 15% proteins and 43% fat; group fed a high-fat diet (HFD, 5.15 kcal/g; Ssniff E15741-34) consisting of 20% carbohydrates, 20% proteins, and 60% fat; and the group fed with a ketogenic diet (KD, 7.55 kcal/g; Ssniff E15149-30) consisting of 6% proteins and 94% fat. The weight of mice and food intake were measured weekly (RADWAG WTC 2000).

### Metabolic cages

After 8 weeks of feeding with the different diet types, six randomly chosen mice from each group were examined in the TSE Phenomaster System to assess metabolic, behavioral, and physiological differences between groups.

Mice spent 10 days in the metabolic cages, including 3 days of adaptation, 3 days of *ad libitum* feeding, 1 day of fasting and 3 days of feeding again (refeeding). During this period, data concerning food and water intake, body mass, voluntary activity, respiratory exchange ratio (RER), and energy expenditure were automatically acquired. The system provides a readout of parameter data every 15 min. Each absolute energy expenditure value was normalized to the animal’s total body weight from the same time point. Normalized energy expenditure values from the refeeding phase were averaged and summed for the final result.

### Plasma and internal organs collection

Subsequently, mice were sacrificed and blood was collected *via* heart puncture in tubes containing heparin sodium salt. Immediately after, the blood was centrifuged for 10 minutes at 10,000 × g and supernatant (plasma) was collected and stored at −80 for further analysis. At the same time selected organs (brain, heart, kidneys, liver, spleen, testicles, epididymis, thymus and thyroid) were isolated, weighed on an analytical scale (RADWAG AS 220.R2) and stored at −80.

### Glucose, BOH and NEFA measurement

Biochemical metabolic parameters such as glucose, ketone bodies and non-esterified fatty acids were measured in collected mouse plasma samples. Fasting and non-fasting glucose levels were checked using a glucometer (Roche, Accu-Chek PERFORMA). To assess ketosis, the level of ketone bodies (β-hydroxybutyric acid) was determined using a Sigma-Aldrich Ketone Body Assay Kit (Sigma-Aldrich, MAK134). To indicate possible insulin-resistance among obese mice the level of non-esterified fatty acids was measured using an ELISA kit (Sigma-Aldrich, EZRMI-13K). Both assays were conducted according to the attached protocol. For colorimetric analysis, an Eppendorf BioSpectrometer basic was used.

### GC-MS analysis

Combined plasma samples were prepared for GC-MS analysis according to the protocol described previously (Wawrzyniak et al., 2018). In brief, 1 µl of 250 mM CaCl_2_ and 1 µl of proteinase K (20 mg/ml) were added to 50 µl of plasma sample and incubated at 37 °C for 60 minutes min. Next, 350 µl of cold methanol:ethanol (1:1, v/v) solution was added, mixed for 5 minutes and stored at −20 °C for 1 hour. Samples were centrifuged at 20 000 x g for 30 minutes at 4 °C. Supernatants were collected and vacuum-dried (SpedVac, Eppendorf). Dry samples were suspended in 50 μl of methoxylamine (15 mg/ml in pyridine) and incubated at 70 °C for 30 minutes. Afterwards, 50 µl of N,O-Bis(trimethylsilyl)trifluoroacetamide with trimethylchlorosilane were added and the mixture was heated again at 70 °C for 30 minutes (Karasinski et al., 2018). Immediately after derivatization samples were subjected to GC-MS analysis as soon as possible. GC-MS analysis was performed using an Agilent 7890 B GC coupled to a quadrupole time-of-flight mass spectrometer 7200 (Agilent Technologies) equipped with an electron impact ionization source. The ZB-5MS column (30 m, 0.25 mm, 0.25 µm, Phenomenex Inc.) was used. The carrier gas was helium at a constant flow rate of 1.2 ml/min. The injection volume was 1 μl with two injection modes - Splitless and Split 30:1. Applied elution program was as follows: the initial temperature of 100 °C was held for 3 minutes, then increased to 290 °C at a constant rate of 5 °C/min and hold for 10 minutes. The total time of chromatographic separation was set to 51 minutes. The temperature of the transfer line and ion source were 300 °C and 280 °C, respectively. Spectra were acquired in the full scan mode applying the m/z range 80–800. The energy of electron ionization was 70 eV.

Raw data was deconvoluted using Unknown Analysis (Version B.09.0, Agilent Inc.) and submitted to statistical analysis using Mass Profiler Professional (Version B.14.5, Agilent Inc.) applying the condition that at least 80% of samples have the analytical signal in at least one of four types of diet plasma samples. Samples were compared pair-wise using an ANOVA test with p≤0.05 and two-fold changes, three and four. Compounds were identified by comparing their mass fragmentation pattern with reference ones with match above 85% using NIST 11 Mass Spectral Library and Human Metabolome Database (HMDB). Principal Component Analysis was performed to find possible relationships among diets.

### miRNA preparation

The pooled plasma samples have been pooled as described above and miRNAs were isolated with mirVana™ PARIS™ Kit (Ambion, USA). All procedures were proceeded according to the manufacturer’s protocols. Briefly, the sample and 2X Denaturing Solution were mixed in equal volumes (100 µl of each) at room temperature (RT), then 200 µl Acid-Phenol: Chloroform was added, and sample was vortexed, centrifuged for 5 minutes at 14000 × g (RT) and aqueous phases were collected to fresh tubes. RNA was eluted with 99% ethanol in a 1:1.25 volume ratio, mixed, and transferred onto a Filter Cartridge followed by centrifugation for 30 seconds with 10000 × g. The sample was washed three times with washing buffers, then 100 µl of preheated (95 °C) nuclease-free water was added and centrifuged for 30 seconds with 10000 × g for RNA collection. Samples were used for reverse transcription reaction with TaqMan MicroRNA Reverse Transcription Kit (Applied Biosystems, USA). Briefly, 3 µl of RNA sample was mixed with RT Reaction Mix, which contained a set of predefined pools of primers for mature miRNAs. PCR reaction was performed in the Mastercycler Nexus Gradient Thermal Cycler (Eppendorf, Germany) with 40 cycles (16 °C for 2 minutes, 42 °C for 1 minute, 50 °C for 1 second), then it was stopped by incubating at 85 °C for 5 minutes. In the next step, a preamplification reaction was prepared – the RT reaction product was mixed with Preamplification Reaction Mix, which contained TaqMan™ PreAmp Master Mix and predefined Megaplex™ PreAmp Primers. The samples were incubated on ice for 5 minutes and a PCR reaction was run with the following settings: 95 °C for 10 minutes, 55 °C for 2 minutes, 72 °C for 2 minutes, and 12 cycles: 95 °C for 15 seconds, 60°C for 4 minutes, then enzyme was inactivated with incubation at 100 °C by 10 minutes. The reaction products were diluted with 75 µl of TE buffer.

### TaqMan arrays

The RT reaction products were mixed with TaqMan™ Universal Master Mix II, no UNG (Applied Biosystems, USA), and loaded to TaqMan™ Array Rodent MicroRNA Card A and B (Applied Biosystems, USA). The card with the sample was centrifuged for 1 minute with 330 × g and run in QuantStudio 12K Flex ™ Real-Time PCR System (Applied Biosystems, USA) with the following reaction parameters: 95 °C for 10 minutes, 40 cycles: 95 °C for 15 seconds, 60 °C for 1 minute. Results were calculated with the ΔΔCt method – data were normalized on global miRNA expression of each sample and mean miRNA expression in samples from STAND group. Data were analyzed with GraphPad Prism 9 (GraphPad Software, USA), and values were presented as the mean of each group ± SD. Statistical analyses were performed using one-way ANOVA, followed by the Tukey’s post-hoc test.

### Gut microbiome analysis

At the end of the 8-week diet, fecal samples were collected in sterile conditions. The samples were stored at −80 °C and sent to GenXone where total DNA was extracted. DNA isolation was performed by using Bead-Beat Micro AX Gravity (A&A Biotechnology). Data from microbiome NGS 16sRNA analysis was assigned to taxa by using the BLAST NCBI 16S Microbial database (day 181007).

To identify microorganisms characteristic to a particular diet Boruta test (Kursa & Rudnicki, 2010) was applied with a standard, normalized Random Forest importance source, using 5000 trees and 200 iterations. For the fast and memory-efficient implementation of Random Forest the R package ranger was used (Wright & Ziegler, 2017). For stability, the Boruta test was performed 50 times and extracted consensus species confirmed in at least 25 iterations.

The diversity of species was quantified by a Shannon diversity index, which is identical to a maximum likelihood Shannon entropy of the distribution of species’ abundances.

### Behavioral tests / Data analysis

In the IntelliCage behavioral test, nosepoke events were collected continuously throughout 9 days in clockwise and 8 days in counter-clockwise tasks, in both active/dark and inactive/light phases. Since some of these tests were carried out at various switch times of the 12h/12 h light cycle, to simplify the analysis timelines of data series resulting from the separate experiments were temporally aligned so that their first active phases begin at the same time. The nosepoke events were then analyzed with a custom Python script according to the following procedure: tracking of an animal began in the active phase when the animal made a lick in the initially expected corner (assigned identically for all animals in a cage). This was counted as the animal’s first correct visit.

Upon any correct visit featured with a lick, the current expected corner assigned to an animal was immediately updated in the clockwise direction. Throughout the test, the animal was allowed to drink (make licks) only in the corner that was its current expected corner. A visit to the current expected corner, with or without a lick, was counted as a correct visit. Incorrect visits was counted when an animal visited a corner which is neither the previous expected corner nor the current expected corner. When an animal made an immediate return (i.e. when the previous expected corner was visited again and no other corner had been visited in the interim), this was counted as a neutral visit, however any further immediate returns were counted as incorrect visits. If prior to revisiting the previous expected corner the animal had made a visit in any non-current expected corner, this was interpreted as a permitted “post-exploratory” revisit and as such was counted as a neutral one. Any other returns were counted as incorrect visits. The ultimate daily proportion of correct visits (% correct visits), shown in Figure 5, was computed as the ratio of the number of correct visits to the sum of the correct and incorrect visits. The number of each type of visit in the CW and CCW tests is illustrated in Supplementary Figure 4.

Data were analyzed with GraphPad Prism 9 (GraphPad Software, USA). In the first analysis (Figure 5, C & D) values were presented as the mean of each group for each day ± CI. Statistical analyses were performed using one-way ANOVA, followed by the Dunnett’s post hoc test and day one was used as a control to which data from the other days were compared. In the second analysis (Figure 5, E & F) % correct visits in day 1, 3, 5 and 7 were presented as mean ± SD and statistical analyses were performed using two-way ANOVA, followed by the Tukey’s post hoc test.

## Aknowledgments

This work was supported by the ANIMOD project within the Team Tech Core Facility Plus program of the Foundation for Polish Science, co-financed by the European Union under the European Regional Development Fund and by National Science Centre (OPUS) grant 2019/35/B/NZ4/02831 (to W.K.).

---

**Supplementary Figure 1. Microbiota diversity and total abundance analysis.** Differences in the fecal microbiome in relation to diet, by phylum. **A,** Shannon diversity, **B,** Total abundance.

**Supplementary Figure 2. Heatmap analysis of gut microbiota changes under the influence of different diets.**

Heat map showing the abundance of species relevant to the diet used, as identified by machine learning analysis (Kursa & Rudnicki, 2010). Abundance is shown as rank for all mice, with blue indicating the lowest and red the highest abundance.

**Supplementary Figure 3. Principal component analysis (PCA) of blood metabolites of mice fed different diets.**

Principal component analysis (PCA) model was obtained for blood metabolites of mice fed different diets (n=4). **A**, graph of results - colors correspond to blood samples taken from mice; orange (STAND) - standard diet, red (HSD) - high sugar diet, green (HFD) - high fat diet, purple (KD) - ketogenic diet; **B,** graph of X loadings shows the distribution of variables in relation to diet types.

**Supplementary Figure 4. Complete data set for different types of visits to diet-dependent cognitive ability tests (clockwise and counterclockwise tests)**

The number of correct (**A, E**), incorrect (**B, F**), neutral (**C, G**), and total (sum of three others) (**D, H**) visits in clockwise (CW) and counter-clockwise (CCW) tests, respectively. Data from each time point were compared with the first day of the experiment for each diet. Experiments were conducted for 9 (CW) or 7 (CCW) days ((CW n: STAND = 46; HFD = 21; HSD = 23; KD = 8); (CCW n: STAND = 19; HFD = 10; HSD = 11; KD = 8), mean ± SD). \**p* ≤ 0.05, \*\**p* ≤ 0.01, \*\*\**p* ≤ 0.001, \*\*\*\**p* ≤ 0.0001; one-way ANOVA with Dunnett’s post hoc test (comparing each time point to week 0, the color of the asterisks is appropriate to the type of diet).

