## Supplementary figures and images for "Hormetic curve of dietary mono- and disaccharide content determines weight gain, gut microbiota composition and cognitive ability in mice"

### Supplementary Figure 1

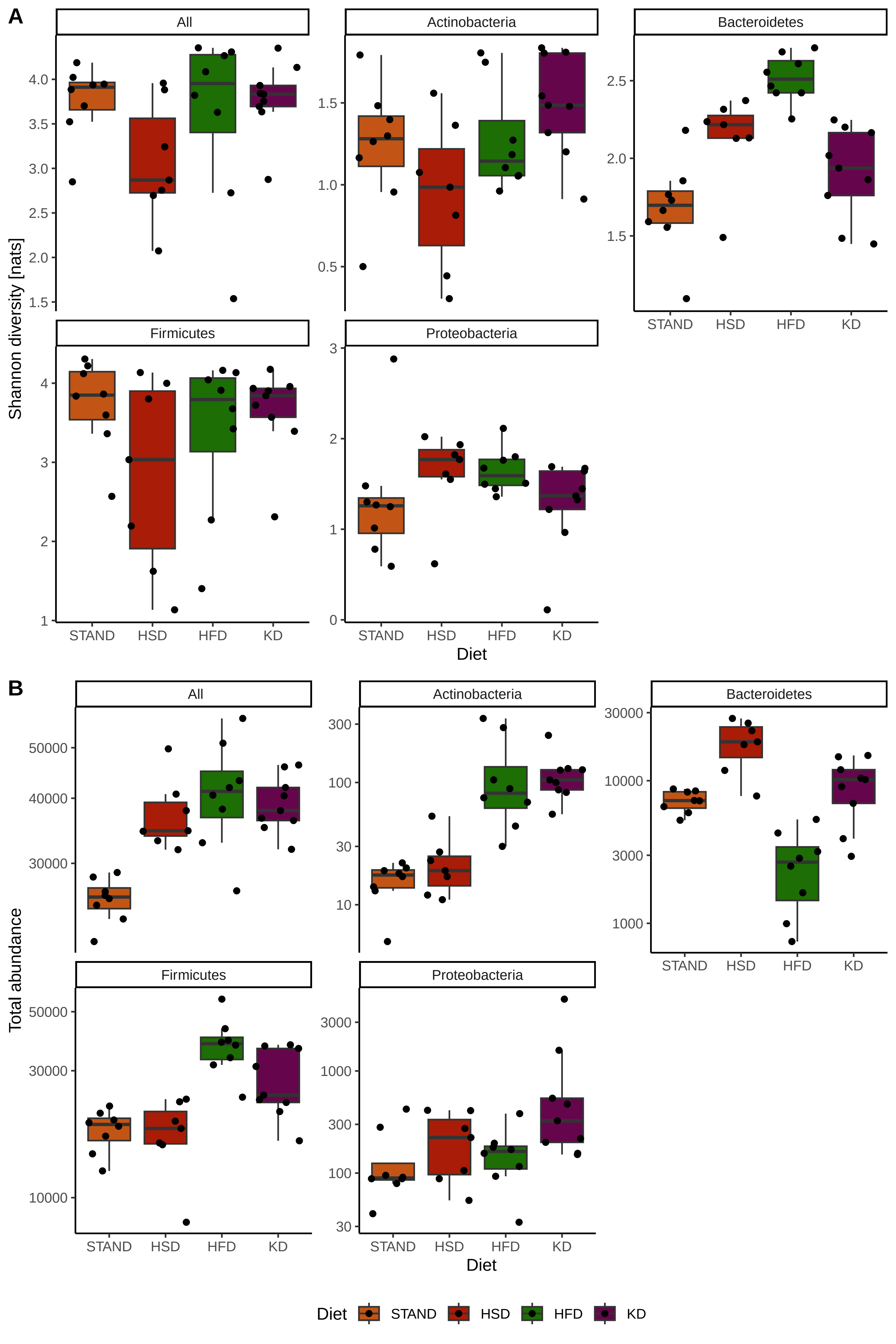

### Supplementary Figure 2

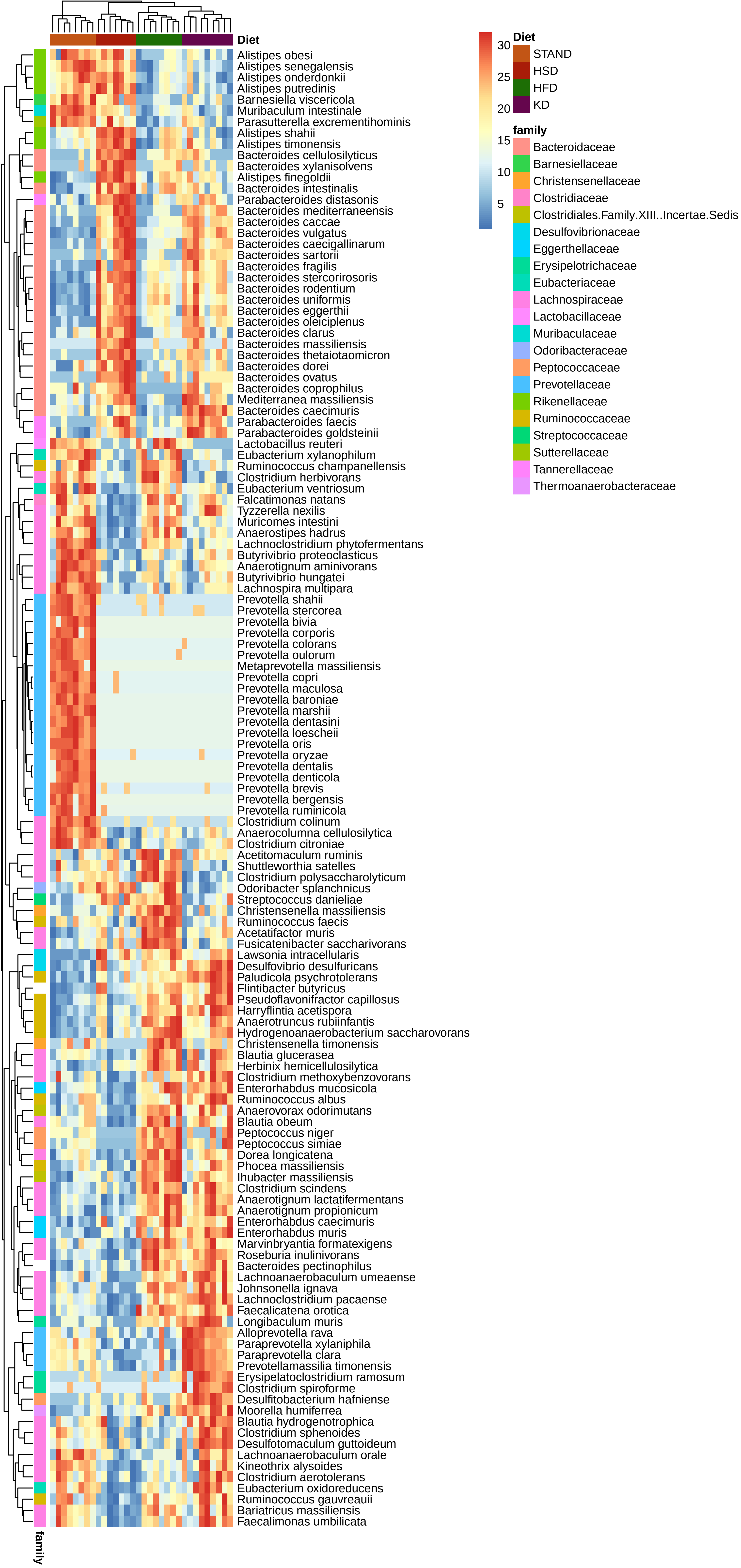

### Supplementary Figure 3

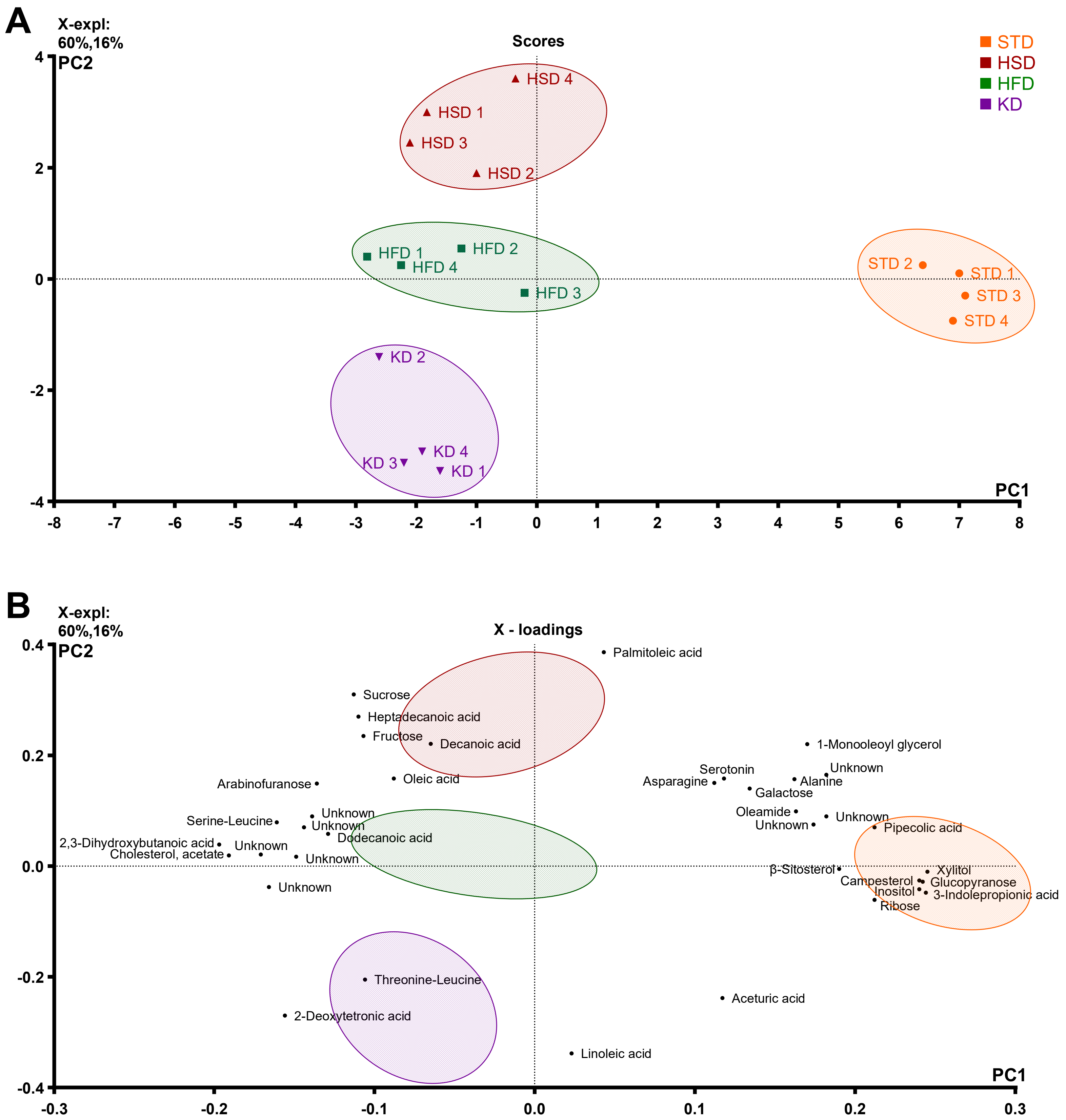

### Supplementary Figure 4

#### CLOCKWISE

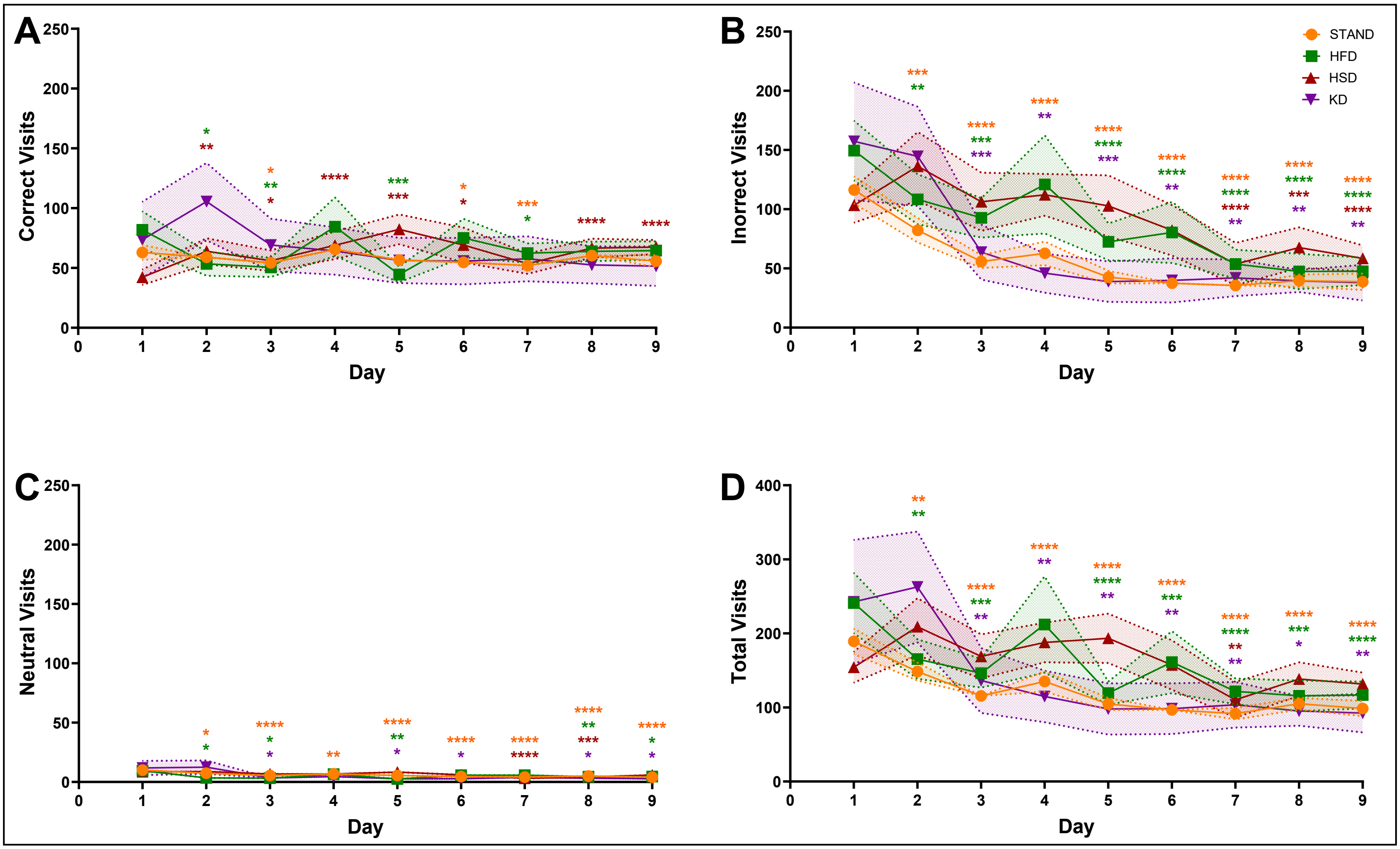


#### COUNTER - CLOCKWISE

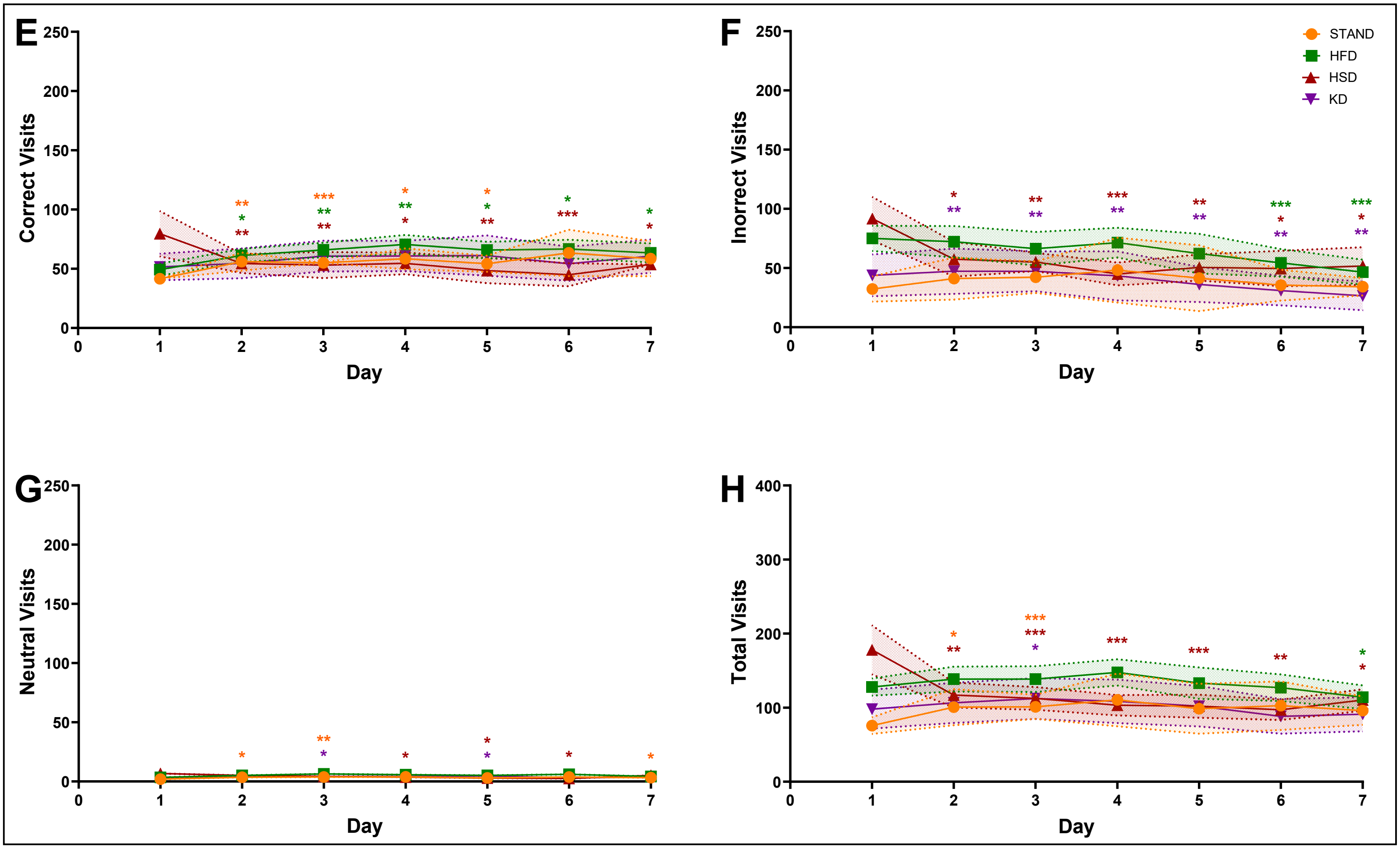
